# Integrating Biomedical Research and Electronic Health Records to Create Knowledge Based Biologically Meaningful Machine Readable Embeddings

**DOI:** 10.1101/540963

**Authors:** Charlotte A. Nelson, Atul J. Butte, Sergio E. Baranzini

## Abstract

In order to advance precision medicine, detailed clinical features ought to be described in a way that leverages current knowledge. Although data collected from biomedical research is expanding at an almost exponential rate, our ability to transform that information into patient care has not kept at pace. A major barrier preventing this transformation is that multi-dimensional data collection and analysis is usually carried out without much understanding of the underlying knowledge structure. In an effort to bridge this gap, Electronic Health Records (EHRs) of individual patients were connected to a heterogeneous knowledge network called Scalable Precision Medicine Oriented Knowledge Engine (SPOKE). Then an unsupervised machine-learning algorithm was used to create Propagated SPOKE Entry Vectors (PSEVs) that encode the importance of each SPOKE node for any code in the EHRs. We argue that these results, alongside the natural integration of PSEVs into any EHR machine-learning platform, provide a key step toward precision medicine.

## INTRODUCTION

The rate at which the ever growing body of world data is being transformed into information and knowledge in some areas (e.g. banking, e-commerce, etc.) far exceeds the pace of such process in the medical sciences. This problem is widely recognized as one of the limiting steps in realizing the paradigm of precision medicine, the application of all available knowledge to solve a medical problem in a single individual (National Research Council, 2011; Colijn et al. 2017).

In order to address this issue, several efforts to integrate these data sources in a single platform are ongoing (Sinha et al, 2015; Chen et al., 2016). The basic premise of data integration is the discovery of new knowledge by virtue of facilitating the navigation from one concept to another, particularly if they do not belong to the same scientific discipline. One of the most promising approaches to this end makes use of heterogeneous networks. Heterogeneous networks are ensembles of connected entities with multiple types of nodes and edges; this particular disposition enables the merging of data from multiple sources, thus creating a continuous graph. The complex nature and interconnectedness of human diseases illustrates the importance of such networks (Barabási et al., 2011). Even bipartite networks, with only two types of nodes, have furthered our understanding on disease-gene relationships, and provided insight into the pathophysiological relationship across multiple diseases (Goh et al., 2007).

In an attempt to address one of the most critical challenges in precision medicine, a handful of recent studies has started to merge basic science level data with phenotypic data encoded in electronic health records (EHRs) to get a deeper understanding of disease pathogenesis and their classification to enable rational and actionable medical decisions. One such project is the Electronic Medical Records and Genomics (eMERGE) Network. The eMERGE consortium collected both DNA and EHRs from patients at multiple sites. eMERGE and subsequent studies showed the advantages of using EHRs in genetic studies (Denny et al., 2010; Ritchie et al., 2010; Kho et al., 2011). Another project linked gene expression measurements and EHRs, an approach through which researchers were able to identify possible biomarkers for maturation and aging (Chen et al., 2008). While these studies illustrate the benefits of combining data from basic science with EHRs, no efforts connecting EHR to a comprehensive knowledge network have been yet reported.

This study builds upon these concepts and utilizes a heterogeneous network called Scalable Precision Medicine Oriented Knowledge Engine (SPOKE) to interpret data stored in electronic health records (EHR) of more than 800,000 individuals at UCSF. Currently, SPOKE integrates data from 29 publicly available databases and contains over 47,000 nodes of 11 types and 2.25 million edges of 24 types, including disease-gene, drug-target, drug-disease, protein-protein, and drug-side effect (Himmelstein and Baranzini, 2015, Himmelstein et al, 2017).

In this work we describe a method for embedding clinical features from EHRs onto SPOKE. By connecting EHRs to SPOKE we are providing real-world context to the network thus enabling the creation of biologically and medically meaningful “barcodes” (i.e. embeddings) for each medical variable that maps onto SPOKE. We show that these barcodes can be used to recover purposely hidden network relationships such as *Disease*-*Gene, Disease-Disease, Compound*-*Gene*, and *Compound*-*Compound*. Furthermore, the correct inference of intentionally deleted edges connecting *SideEffect* to *Anatomy* nodes in SPOKE is also demonstrated.

## RESULTS

The main strategy of this work is to embed EHRs onto the SPOKE knowledge network utilizing a modified version of PageRank, the well-established random walk algorithm (Page et al., 1999). These embeddings, called Propagated SPOKE Entry Vectors (PSEVs) encode the importance of each node in SPOKE for every overlapping concept between the EHRs and SPOKE.

### Embedding EHR concepts in a knowledge network

Deidentified EHR data from 816,504 patients was obtained from the UCSF medical center through the Bakar Computational Health Sciences Institute (BCHSI). The cohort was then filtered to only include patients that had been diagnosed with at least one of the 137 complex diseases currently represented in SPOKE, leaving 292,753 patients for further analysis. Select structured data tables (including medication orders, lab tests, and diagnoses) were used to create the PSEVs (see Methods). Each structured EHR table contains codes that can be linked to standardized medical terminology allowing direct links to SPOKE, referred to as SPOKE Entry Points (SEPs). There are currently 3,527 SEPs and although this represents a sizable proportion (7.5%) of all nodes in SPOKE, most nodes are not directly reachable, thus potentially diluting the power of the network’s internal connectivity. Thus, a modified version of the random walk algorithm was used to propagate SEPs through the entirety of the knowledge network, thus creating a unique medical profile for each of the selected clinical features in the EHRs.

In the original random walk algorithm, a walker is placed onto a given node in a network, and it can move from one node to another as long as there is an edge connecting them. The algorithm was adjusted in a way similar to topic-sensitive PageRank (Haveliwala 2002), by weighing the re-start parameter of the random walker towards nodes that are important for a given patient population.

This modified version of PageRank can be applied to any patient cohort. To demonstrate that these vectors capture biologically meaningful information, PSEVs for Body Mass Index (BMI) (an ubiquitous variable in the EHR) were created. A patient’s BMI is recorded at each visit and is equivalent to their weight (kg) over their height (m) squared. BMI is typically used to classify patients into 4 main categories (underweight, normal, overweight, and obese). Decades of research have provided deep insight into both the phenotypic and mechanistic manifestation of obesity. However, only the top-level (phenotypic) information (i.e. BMI category) is captured in the EHRs. We hypothesized that by using this method it would be possible to integrate mechanistic and biological level data.

When examining the distribution of BMIs across the UCSF patient population four groups are clearly distinguishable. Though these four groups align well with the standard categories, patients were separated in unbiased manner using k-means clustering in order to keep the algorithm blind to these pre-assigned classes (Figure 1A). Therefore, patient cohorts can be created without a priori knowledge of the standard classes. Figure 1B illustrates the modified PageRank algorithm using patients in the high BMI cohort (BMI > 34). First, the records from all 73,237 patients in the high BMI cohort were extracted. Second, connections were created between each of those patients and all of their additional SEPs. By definition, this means connections to any medication, diagnosis, or laboratory result that is present in both that patient’s record and SPOKE. Third, a random walker is initialized and allowed to randomly jump back to the patient population with probability β (optimized β=0.1). Each iteration results in a rank vector that reflects the proportion of time the walker spends on each node in the network. In practice, for each iteration, this is calculated by taking the dot product of the transition probability matrix and the rank vector from the previous iteration (See Methods). Once the difference between the previous and current rank vector is less than some threshold (alpha=1E-3), the final PSEV is returned (bottom vector).

**Figure 1:**
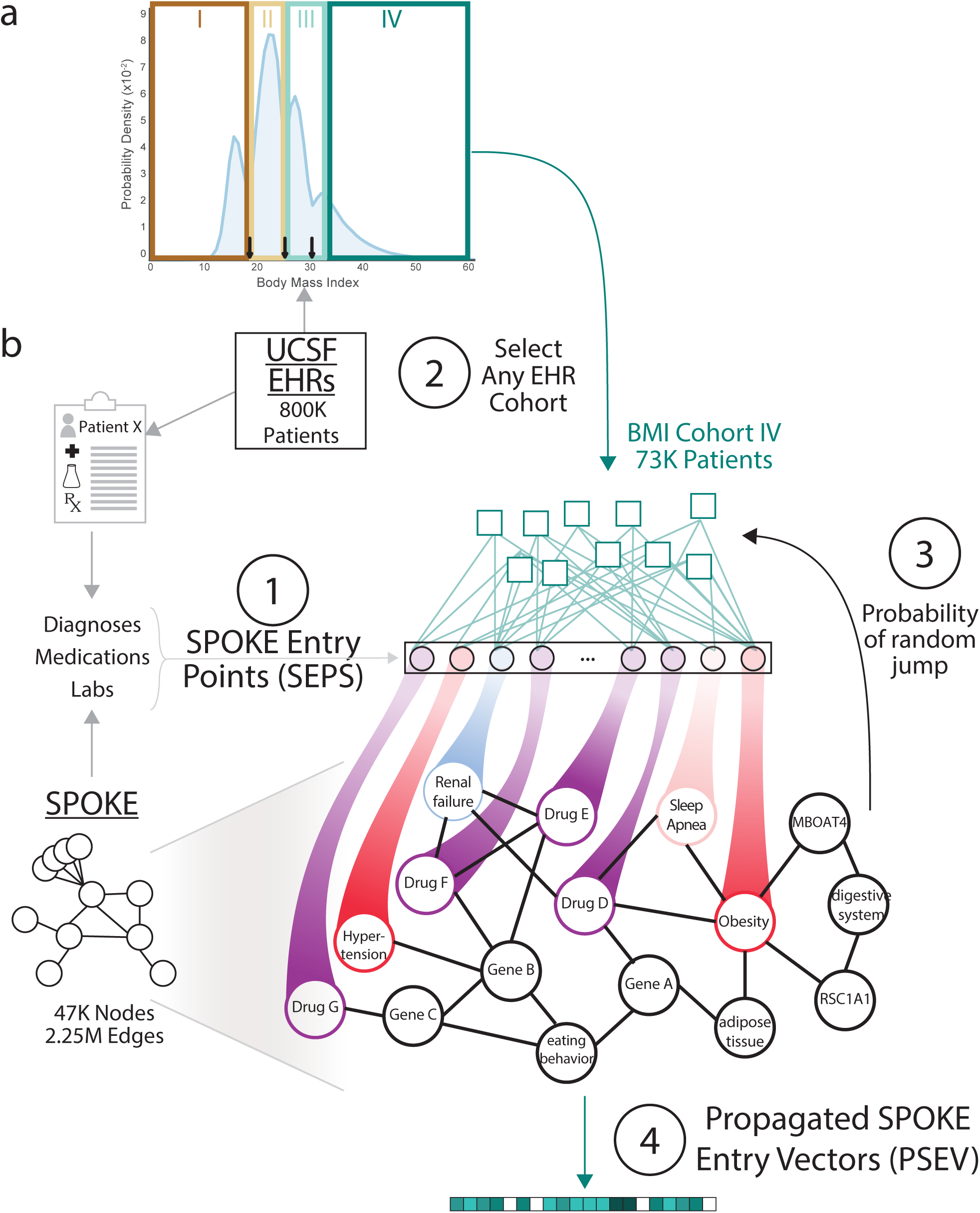
Embedding EHR concepts in a knowledge network. **(a)** Distribution of patient BMIs at UCSF. Four BMI cohorts were created using K-means clustering of BMI (boxes I-IV : <=19, 19.1-25.5, 25.6-34.2, and >34.2). Arrows at the bottom correspond to the BMIs that separate the standardize weight classes. **(b)** Step 1: find the overlapping concepts between SPOKE and the patient data (EHRs). These are called SPOKE Entry Points (SEPs). Step 2: choose any code or concept in the EHR to make cohort. Here we have chosen patients with a high BMI (Cohort IV). Then connect each patient in the cohort to all of the SEPs in their records. Step 3: perform PageRank such that the walk-er restarts in the patient cohort. Iterate until desired threshold is reached. Step 4: final node ranks are then used to create the weights in the Propagated SPOKE Entry Vector (PSEV).

One can imagine SPOKE as a set of interconnected water pipes and the SEPs as input valves. Then, the percentage of high BMI patients that have type 2 diabetes in their EHRs will determine how much water is allowed to flow through the type 2 diabetes SEP “valve”. Once all of the valves have been calibrated to fit the high BMI patient population, water can then flow to downstream nodes in SPOKE. Once the water reaches an almost steady state, differential water flow will highlight intersections of pipes (SPOKE nodes) that are significant for high BMI patients.

### Identifying Phenotypic Traits in PSEVs

The final PSEV is representative of how important each SPOKE node is for a given EHR concept based on both the connections in SPOKE and the patients with that concept in their EHR. To examine what this means, the prioritization of *Disease* elements in the PSEVs were compared for each of the four BMI cohorts. The top *Diseases* in the PSEV of the highest BMI cohort are obesity, hypertension, type 2 diabetes mellitus, and metabolic syndrome X. While not unexpected, the identification of these diseases as the most important conditions for this group of patients without any reference to the mechanisms underlying obesity present in the EHR, is noteworthy. These diseases are also well correlated with average BMI (r=0.75-0.95) and when their rank is plotted against average BMI, have some of the steepest slopes (slope=5.4-6.7), suggesting they are causally related.

To learn more about the relationships between BMI and these diseases the plots of rank vs average BMI were further examined (Figure 2A). Hypertension becomes a top ranked *Disease* almost immediately (moving from rank 133 to 6 between BMI categories I and II). This makes sense given that hypertension is the most prevalent disease in UCSF cohort and many of the factors that contribute to hypertension risk are also related to increasing BMI. Metabolic syndrome X and obesity also display an abrupt rank change, on average 128 positions, between BMI categories II and III. This change suggests that metabolic syndrome X and obesity become associated with BMI once people have reached overweight status and that an increased BMI is one phenotypic manifestation of these conditions. Finally, type 2 diabetes mellitus becomes significantly ranked (position 4) when patients reach overweight status. However, it differs in that progression in rank between BMI categories I and III is gradual suggesting increased BMI as a risk factor in type 2 diabetes mellitus. In contrast, celiac and Crohn’s disease progressively move down 114 and 120 positions respectively between BMI categories I and IV. This trend could be explained by the fact that weight loss is symptomatic of both celiac and Crohn’s disease. Another *Disease* that shows a progressively moves down in rank with increased BMI is attention deficit hyperactivity disorder (ADHD). This negative correlation is due to the fact that most of the medications used to treat ADHD have side effects related to weight loss and loss of appetite. These results show that the algorithm correctly up-weights phenotypes associated with high BMI in the PSEVs for Cohorts III and IV while also down-weighting those phenotypes in the low BMI Cohorts.

**Figure 2:**
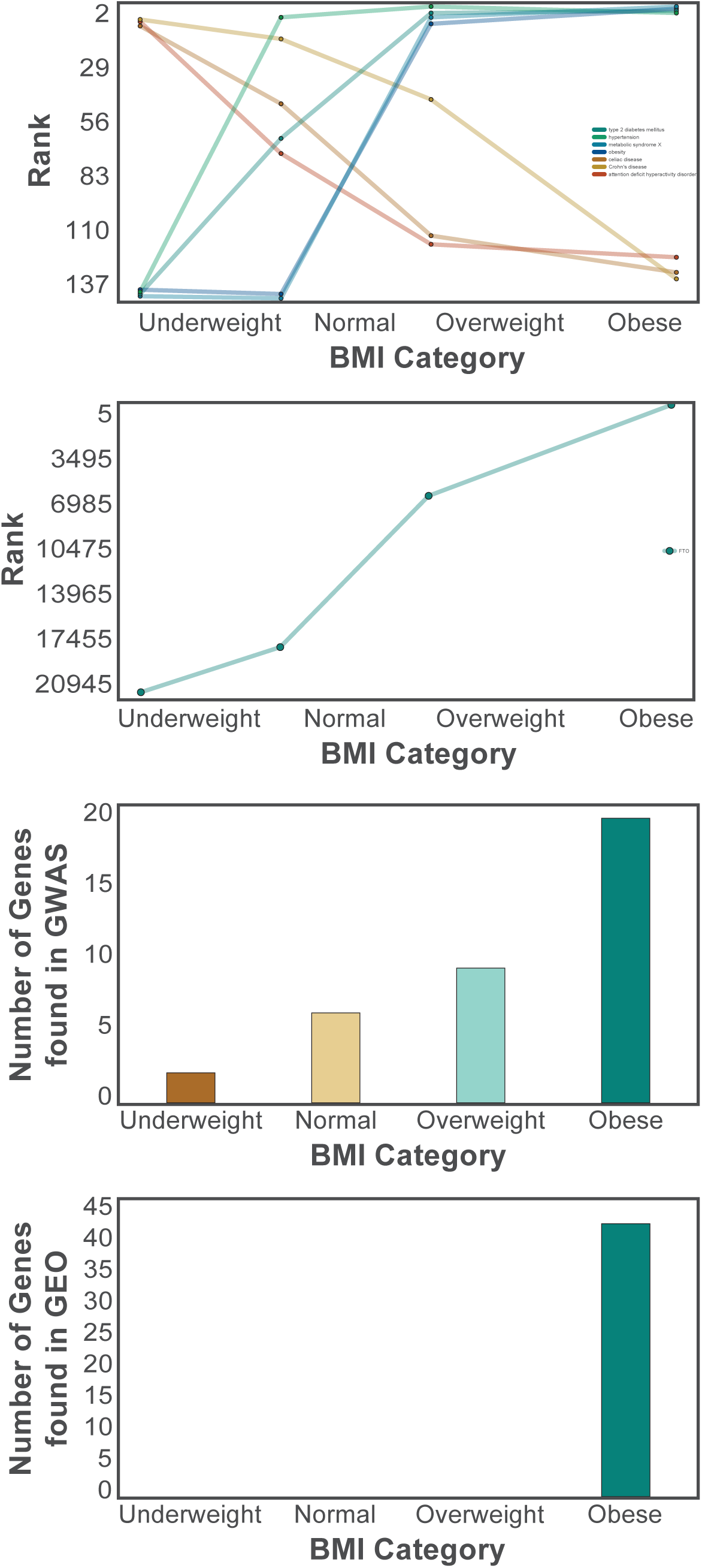
PSEVs contain phenotypic and genotypic information. **(a)** BMI Cohort vs Disease Rank. The top 4 ranked Diseases in the in Cohort IV’s PSEV are obesity, hyperten-sion, type 2 diabetes mellitus, and metabolic syndrome X. All 4 show a positive relationship with BMI. The opposite trend is observed for celiac disease, Crohn’s disease, and attention deficit disorder which are highly ranked in Cohort I’s PSEV. **(b)** FTO gene is positively correlated with BMI. **(c)**The number of over-lapping genes between the GWAS catalog for increased BMI and the top 2.5% of Genes in each BMI cohort PSEV. **(d)** The number of overlapping genes between BMI related GEO datasets and the top 2.5% of Genes in each BMI cohort PSEV.

It should be noted that up until this point, BMI has been treated as a continuous variable used to simply split patients into groups and the algorithm has been blind to the standardized classes associated to those groups. BMI was chosen to illustrate the utility of PSEVs because the consequences/traits of an abnormal BMI are very well known. However, since a PSEV can be created for any variable in the EHRs they can also be used to reveal phenotypic traits associated with less well-understood variables and phenotypes.

### PSEVs can Learn Genotypic Traits and Underlying Biological Mechanisms

To test whether the same trend was seen at genotypic level, linear regressions were computed on the average BMI vs *Gene* rank. Again, the genes that positively correlated with average BMI were given the top prioritization in the high BMI PSEV. An example of a gene that is positively correlated with BMI is Alpha-Ketoglutarate Dependent Dioxygenase (FTO), also known as Fat Mass And Obesity-Associated Protein, is shown in Figure 2B. To check if these genes were genetically related to BMI, genes associated with increased BMI (not necessarily obesity, just an average increase) were extracted from the GWAS catalog (n=365) and compared them to the top 365 ranked Genes in the PSEVs. Remarkably, BMI category IV was significantly enriched in known BMI associated genes (p=2.19E-10; Figure 2C). BMI category III was also significant while the BMI cohorts corresponding to underweight and normal BMIs showed no significant enrichment. Additionally, it was hypothesized that genes with altered expression would also be highly ranked. We found that 34% of dysregulated genes resided in the top 0.6% (n=119) of genes in the PSEV for cohort IV (p=9.28E-72; Figure 2D). This immense enrichment occurred because, unlike the GWAS catalog, datasets in the Gene Expression Omnibus (GEO) with just BMI as a phenotype (without any other major disease), had already been incorporated into SPOKE via obesity *Disease*-UP(DOWN)REGULATES-*Gene*. Together these results illustrate that PSEVs can learn new relationships (GWAS) while also maintaining the known relationships in SPOKE (GEO).

### PSEVs Preserve Original SPOKE Edges

After identifying that the high BMI PSEV was able to preserve the known gene expression edges in SPOKE, we decided to check this in a high throughput manner. To do this, PSEVs were created for all of the concepts in the EHRs that directly mapped to a node in SPOKE (SEPs; n=3,233). Then the top ranked nodes (ranked per type) in each PSEV were examined (Supplementary Figure 1A-C). The majority of top ranked nodes in a given PSEV are also first neighbor relationships in SPOKE. For example, the Multiple Sclerosis (MS) *Disease* node is connected to 39 *Anatomy* nodes in SPOKE, if the top 39 ranked *Anatomy* nodes are selected from the MS PSEV there is a 100% overlap with the MS *Anatomy* neighbors. Similarly, for *Symptom* nodes connected to MS, 80% of first neighbor relationships are maintained. This means that although most of the top nodes are the same, new relationships are prioritized based on the symptoms experienced by individual MS patients at UCSF. Next, the prioritizations of nodes that are not directly connected in SPOKE were considered (Supplementary Figure 1C). For instance, multiple nodes related to the *response to interleukin-7* are ranked among the top 10 *BiologicalProcess* nodes and the node for the *structural constituent of myelin sheath* in the top 10 *MolecularFunction* nodes. Though there is an abundance of evidence supporting these relationships, there is no direct relationship in SPOKE nor is this information stored in the EHRs, thus they must be learned during PSEV creation. These results illustrate the ability of PSEVs to preserve the original information from SPOKE while expanding its significance in a biologically meaningful manner by reaching out to more distant but biologically related nodes. Further, this demonstrates that PSEVs describe each EHR concept in multiple dimensions and is true to the hierarchical organization of complex organisms.

After identifying and implementing a method to embed EHR onto the knowledge network, we sought to verify in a rigorous manner that the obtained vectors are biologically meaningful (i.e. that the expanded set of variables stemming from EHRs result in a network of related medical concepts). Next, we demonstrate that the PSEV ability to learn genetic relationships can be applied in a high throughput fashion.

Additionally, a series of benchmarks (supplemental text) shows that PSEVs ability to learn connections can be applied to other edge types such as *Disease-Disease* and *Compound-Compound* similarity, *Compound* to drug-protein (molecular targets), and *SideEffect-Anatomy*.

### Uncovering specific *Disease*-*Gene* Relationships in EHR embeddings

Because of the multitude of concepts present in SPOKE, multiple paths can connect any two nodes, thus providing redundancy. Thus, we hypothesized that unknown relationships, like the GWAS genes recovered in the high BMI PSEV, could still be inferred even if some of the information was missing because the random walker would traverse similar paths during PSEV computation. To address this point, all of the *Disease*-*Disease* and *Disease*-*Gene* edges in SPOKE were removed and the PSEVs were recomputed the *Disease* PSEVs (PSEV^ΔDD, ΔDG^), ranking the *Gene* nodes in each *Disease* PSEV.

The resulting PSEVs (PSEV^ΔDD, ΔDG^) were visualized in a heatmap and clustered by *Diseases* and *Genes* (Fig 3A). Clearly defined groups of diseases can be identified in the heatmap, many of which are known to share associated or influential genes. For example, Disease Cluster 4 contains mainly neurological, diseases such as multiple sclerosis, Alzheimer’s disease, narcolepsy, autistic disorder, and attention deficit hyperactivity disorder. The Gene cluster most characteristic of Disease Cluster 4 contains 197 genes (Fig 3B). Within this Gene cluster, 96 *Genes* are associated with at least one *Disease* in Disease Cluster 4 (enrichment fold change=2.0), 33 *Genes* are associated with at least 2 diseases (enrichment fold change=3.9), and 15 *Genes* are associated with at least 3 diseases (enrichment fold change=5.4; Fig 3C-D). These results support the hypothesis that PSEVs encode deep biological meaning.

**Figure 3.**
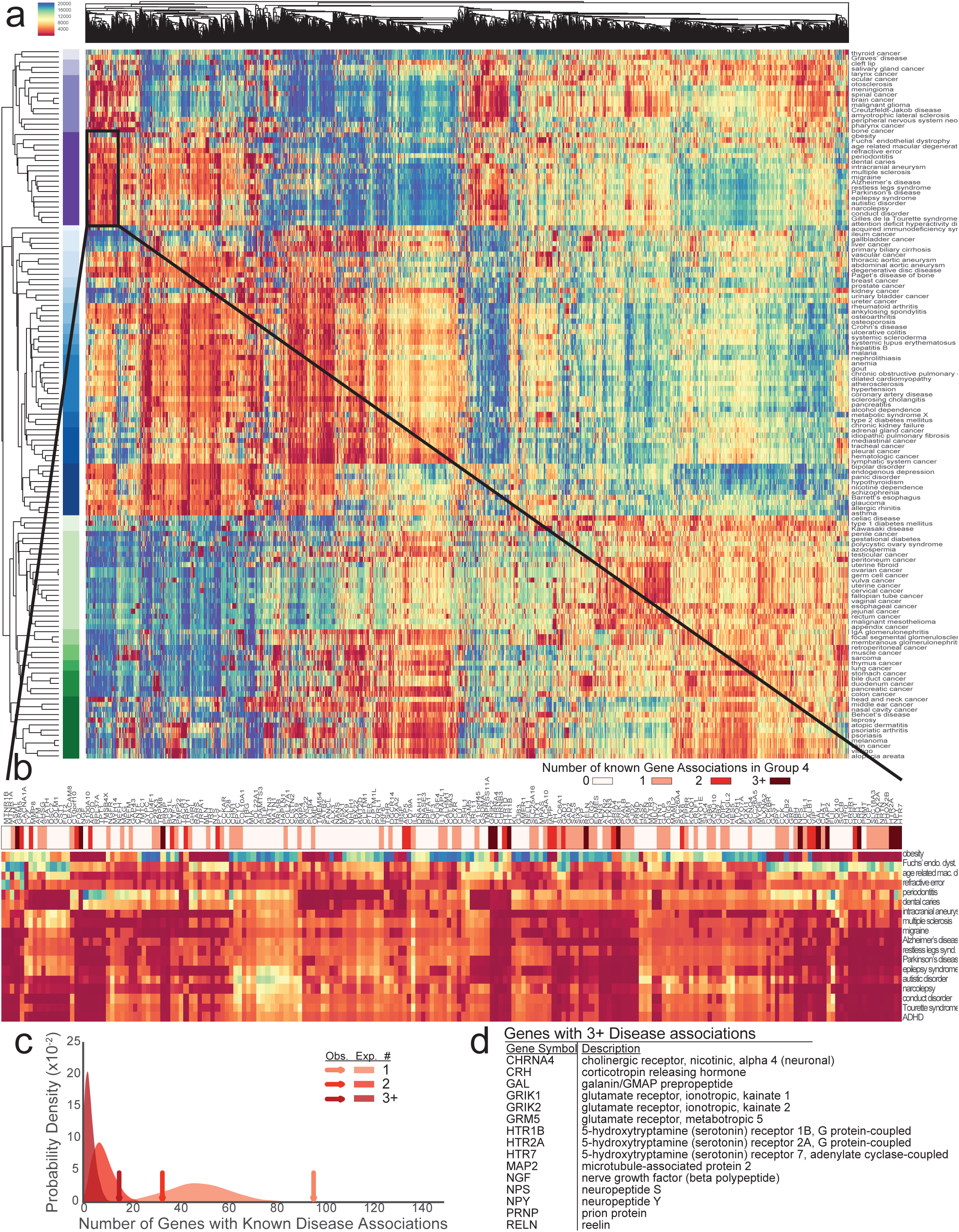
Disease Cluster by Genetic Similarity. **(A)** Heat map generated with the *Disease* PSEV^ΔDD, ΔDG^ (only using elements of *Genes* that associate with at least one *Disease*). Both *Disease* (rows) and Genes (columns) are clustered. Disease Cluster 4 (n=18) is enriched in neurological diseases and shown in dark purple. **(B)** Magnification of the 197 *Genes* found in a top *Gene* Cluster (Cluster 6) for Disease Cluster 4. Asterisks above Gene symbols indicate how many *Disease* in Dis-ease Cluster 4 are associated with that *Gene*. Color bar signifies how many Diseases were associat-ed with a given Gene. **(C)** Expected distributions for the number of *Genes* that are associated with at least one, two, or three Diseases (1000 random permutations of 18 *Disease* and 197 Genes). Arrows show the observed number over *Genes* within Gene Cluster 6 that are associated with at least one, two, or three *Disease* in Disease Cluster 4 and greatly exceed the expected number of *Genes* (fold change=2.0, 3.9, and 5.4 accordingly). **(D)** 15 *Genes* that are within Gene Cluster 6 are associated with three or more *Disease* in Disease Cluster 4.

To validate that the recomputed PSEVs (generated without the critical edges) were able to uncover genetic relationships among the complex diseases in SPOKE, a *Disease*-*Gene* networks (DG) using the top K *Gene* nodes for each *Disease* in PSEV^ΔDD, ΔDG^ was created, where K is equal to the number of known gene associations for a given disease. In SPOKE, the ASSOCIATES_DaG edges represent known associations between *Diseases* and *Genes* and are obtained from the GWAS Catalog (MacArthur et al., 2017), DISEASES (Pletscher-Frankild et al., 2015), DisGeNET (Pin ero et al., 2015; Pin ero et al., 2016), and DOAF (Xu et al., 2012). DG networks were generated using either the original PSEVs (DG^PSEV^, Blue) or the incomplete, benchmarking PSEV^ΔDD, ΔDG^ (DG ^PSEV^^ΔDD, ΔDG^, Green Fig. 4A). These networks were compared against networks created using three random matrices as a way to generate a null distribution: PSEV^RANDOM^ (DG^RANDOM^, Pink distribution Fig. 4A), PSEV^SPOKE SHUFFLED^ (DG^SPOKE^^SHUFFLE^, Red), and PSEV^SEP SHUFFLED^ (Orange, DG^SEP SHUFFLE^). Next, the number of overlapping edges between each of the DG networks and the gold standard *Disease*-ASSOCIATES_DaG-*Gene* (DG^SPOKE^) edges (n=12,623) in SPOKE were compared. When selecting the top K *Genes* using only *Genes* with at least one ASSOCIATES_DaG edge (n=5,392), both DG^PSEV^ and DG^PSEV^^ΔDD, ΔDG^ shared significantly more edges with DG^SPOKE^ than with any of the random networks (Fig 4A; average fold change 15.2 and 2.4 accordingly). This suggests that redundancy in spoke paths can be used to infer genetic relationships even when the original (direct) associations are removed.

**Figure 4.**
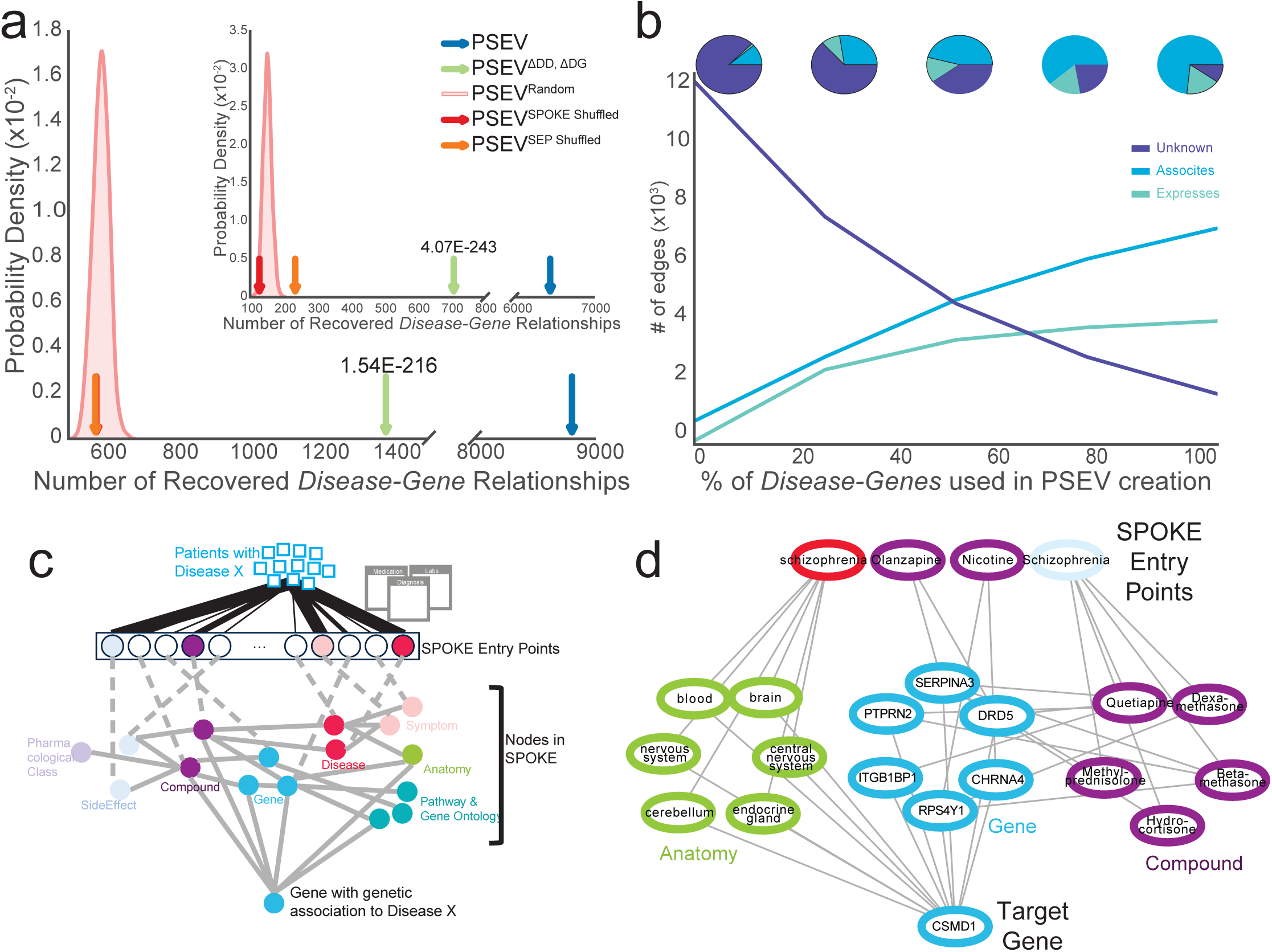
Recovering deleted *Disease-Gene* edges. Prior to PSEV^ΔDD, ΔDG^ calculation all of the *Disease-Gene* and *Disease-Disease* edges were deleted from SPOKE. (A) The gold standard *Disease-Gene* network was made from the deleted edges in SPOKE. Plots show the number of *Disease-Gene* relationships using each of the PSEV matrices that overlap with the gold standard networks. The pink distributions show the results from the permuted PSEV matrices (PSEV^Random^; 1000 iterations) while the arrows show the results from the original PSEV (blue), PSEV^ΔDD, ΔDG^ (green), PSEV^SPOKE SHU^^FFLED^ (red), and PSEV^SEP SHUFFLED^ (orange). **(A)** The top K *Genes* where selected from the set of Genes in the gold standard network or **(A insert)** the entire set of *Gene* nodes in SPOKE. **(B)** The breakdown of top *Disease-Gene* relationships as knowledge (edges) is added back to the net-work. **(C)** To uncover how the deleted *Disease-Gene* associations are recovered using the PSEVs we retraced the shortest path between the most important SPOKE Entry points (SEPs) and the de-sired *Gene*. Patients with *Disease* X put pressure on the SEPs (top). The SEPs that receive the most significant amount of pressure are colored by node type. Information then flows through other nodes in SPOKE (middle) before reaching the Gene that is genetically associated to *Disease* X (bottom). **(D)** In the GWAS catalog Schizophrenia and CSMD1 are associated. As outlined in B, the information flows from the significant SEPs of patients with Schizophrenia to CSMD1.

**Figure 5:**
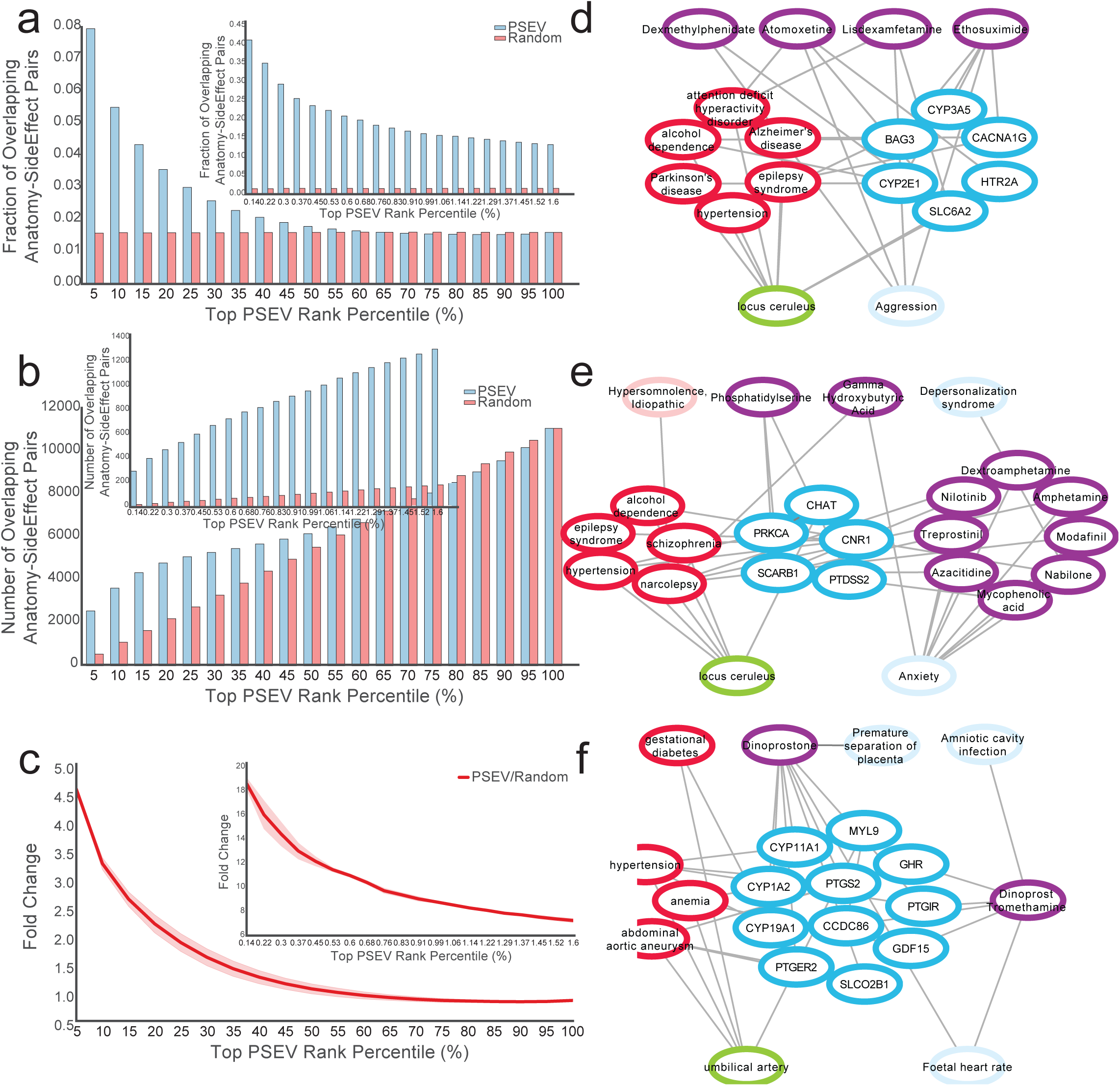
MEDLINE *Anatomy-SideEffect* Relationships are Top Ranked Nodes in PSEV. Fraction **(A)**, count **(B)**, and fold change **(C)** of overlapping edges MEDLINE *Anatomy-SideEffect* network and PSEV *Anatomy-SideEffect* network (blue) or random PSEV *Anatomy-SideEffect* network (red) for different thresholds of PSEV disease similarity. **A-C** Are shown in 5% similarity intervals of ranked nodes starting with the most similar 5% left and all nodes (100%) right. The inserts in **A-C** focus on the top 0.14-1.6% of ranked nodes. **D-F** Examples shortest paths connecting the nodes that contrib-ute the most to the *Anatomy-SideEffect* similarity to the target *SideEffect* and *Anatomy* nodes.

These results were even more striking when selecting the top K genes using all genes in SPOKE (Fig 4A insert; n=20,945; average fold change 40.6 and 4.5 accordingly). It should also be noted that, unlike PSEV^ΔDD, ΔDG^, both PSEV^SEP SHUFFLED^ and PSEV^SPOKE SHUFFLED^ were created without deleting the *Disease*-*Disease* and *Disease*-*Gene* edges from SPOKE, therefore the correct edges were present at least some of the time even in the permuted networks, thus providing a higher level of stringency.

### Learning Rate Differs Between Edge Types

One of the main challenges with knowledge networks is that as long as our knowledge is incomplete, the networks will suffer from missing edges. The benchmark shown here illustrates the most severe scenario in which 100% of our knowledge about the relationships among *Diseases* and between *Diseases* and *Genes* is removed. To evaluate performance of the algorithm as the network gains knowledge, edges were slowly added back to the network. We found that the PSEVs learned well-established (ASSOCIATES) *Disease-Gene* edges before the more noisy (REGULATES) edges (Figure 4B). This is most likely due to the fact that well-established (associated) *Genes* are necessarily drivers of (not reacting to) a *Disease*. In practice this would cause the random walker keep going back to *BiologicalProcess, CellularComponent, MolecularFunction*, and *Pathway* nodes that are important for a given *Disease* and thereby push information to *Genes* involved in those activities. Alternatively, the random walker could travel to *Anatomy* nodes that express *Genes* that are associated with a *Disease* or through *Compounds* that are used to treat (or even those that exacerbate) a *Disease*. This further demonstrates that the relationships inferred within PSEVs are biologically meaningful.

### Retracing the path between SEP and Genes

Finally, to understand how the patient population at UCSF influenced the PSEVs to correctly rank *Disease-Gene* associations the shortest paths were retraced between the significant SPOKE Entry points of a given *Disease* and the associated *Gene* (Fig 4C; Methods). For example, the locus containing *CSMD1* is associated with Schizophrenia in the GWAS Catalog. Figure 4D shows why the gene *CSMD1* was one of the top ranked Genes in the PSEV^ΔDD, ΔDG^ for Schizophrenia. The weight from the EHRs of Schizophrenia patients at UCSF drives information towards *Anatomy* in which *CSMD1* is expressed or regulated and *Compounds* that bind or regulate *Genes* that interact or regulate with *CSMD1*. The combined weight highlights *CSMD1* as a gene that is associated with Schizophrenia. This example highlights the fact that inferences made with this method are not “black box” predictions, but the information used to make the inference can be traced back to the exact concepts. We believe that knowledge based “clear box” algorithms, such as the one presented here, will be pivotal in the advancement of precision medicine.

## DISCUSSION

Uncovering how different biomedical entities are related to each other is essential for speeding up the transformation between basic research and patient care. When deciding the best therapeutic management strategy for a patient, physicians often need to think about the symptoms they present, their internal biochemistry, and potential molecular impact and adverse events of drugs simultaneously. A well-trained and experienced doctor will likely prescribe the best course of action for that patient. However, significant heterogeneity is seen even across the best hospitals on what “best course of action” means for a given patient, resulting in poor consistency, a labyrinth of solutions, and ultimately lack of evidence-based medicine. Since it is naturally impossible for a single person to retain and recall all the necessary and relevant information, an efficient manner to incorporate this knowledge into the health care system is needed. We argue that since PSEVs can be created for any code or concept in the EHRs it is possible they could provide such solution. Using PSEVs we were able to integrate what we have learned from the last five decades of biomedical research into the codes used to describe patients in the EHRs. As a result, these embeddings serve as a first step to bridging the divide between basic science and patient data.

Our method for the integration of EHRs and a comprehensive biomedical knowledge network is based on random walk. Random walk has been applied to a wide variety of biological topics such as protein-protein interaction networks (Can et al., 2005), gene enrichment analysis (Subramanian et al., 2005), and ranking disease genes (Köhler et al., 2008; Valentini et al., 2014; Wang et al., 2015). Additionally, random walk has been used to infer missing relationships in large incomplete knowledge bases (Lao et al., 2011). Our method includes the generation of PSEVs, as a way to embed medical concepts onto the network. The entire patient population at UCSF was used to determine how important each node in SPOKE is for a particular code. Therefore, each PSEV describes EHR codes in both a high level phenotypic and deeper biological manner.

We demonstrate that not only do PSEVs carry the original relationships in SPOKE, but also are able to infer new connections. This was illustrated by ability of PSEVs to recover deleted *Disease-Disease, Disease-Gene, Compound-Compound*, and *Compound-Gene* edges as well as to infer new relationships between *SideEffect* and *Anatomy* nodes. Other than just showing that PSEVs can learn relationships between different types of nodes, these tests illustrated that PSEVs can learn relationships between nodes at a variety of lengths apart from one another. By inferring the *Disease*-*Disease* and *Compound*-*Compound* edges, we demonstrated that PSEVs could find SEP-, or EHR-level relationships. By inferring *Disease*-*Gene* and *Compound*-*Gene* edges, we verified that PSEVs could find SEP to SPOKE level relationships. Finally, by inferring *SideEffect*-*Anatomy* edges we proved PSEVs could find SPOKE-level relationships. These tests served as our proof of principle that PSEVs can learn multiple types of new relationships.

Further, these results illustrate that, unlike black box methods, PSEVs are capable of embedding phenotypic traits such as risks, co-morbidities, and symptoms. Other vectorization methods like word2vec are able to learn relationships, however since the elements within the vector are unknown they cannot be traced back to a given trait in the EHRs. Similarly, though it is possible to identify these phenotypic traits using a statistical analysis of a single cohort, the benefit to using PSEVs is that these traits are identified in a high throughput fashion for every concept in the EHRs and outputs them in a format that can be used in machine learning platforms. PSEVs, and other clear box algorithms, allow us to integrate knowledge into data, therefore generating deeper, informed characterizations that can be understood by both humans and machines.

The potential uses of PSEVs are vast. We recognize that several associations in EHRs can be uncovered using clinical features alone, and several machine-learning approaches are already being utilized to that end (Shickel et al., 2018). However, since PSEVs describe clinical features on a deeper biological level, they can be used to explain why the association is occurring in terms of Genes, Pathways, or any other nodes in a large knowledge network like SPOKE. Consequently, PSEVs can be paired with machine learning to discover new disease biomarkers, characterize patients, and drug repurposing. With implementation of some of these features, we anticipate that PSEVs or similar methods will constitute a critical tool in advancing precision medicine.

## MATERIALS AND METHODS

### Electronic Health Records

The University of California, San Francisco (UCSF) supplied the Electronic Health Records (EHRs) in this paper through the Bakar Computational Health Sciences Institute. Almost one million people visited UCSF between 2011-2017. Out of 878,479 patients 292,753 had at least one of the 137 complex diseases currently represented in SPOKE. The EHRs were de-identified to protect patients’ privacy. For this paper we collected the information on the cohort of patients with complex diseases using de-identified LAB, MEDICATION_ORDERS, and DIAGNOSES tables. The LAB table contains the lab test orders and results, including the actual measurements and the judgment of whether the results were abnormal. The MEDICATION_ORDERS table contains prescriptions with dose, duration, and unit. The DIAGNOSES table contains diagnosis and symptoms with ICD9 and ICD10 codes. These tables are linked by Patient IDs (one unique ID for each patient) and Encounter IDs (one unique ID for each encounter a given patient has with our medical system).

### Scalable Precision Medicine Oriented Knowledge Engine

Scalable PrecisiOn Medicine Knowledge Engine (SPOKE) is a heterogeneous knowledge network that includes data from 29 publicly available databases, representing a significant proportion of information gathered over five decades of biomedical research (Himmelstein et al. 2017). This paper was powered by the first version of SPOKE, which contains over 47,000 nodes of 11 types and 2.25 million edges of 24 types. The nodes (Anatomy, BiologicalProcess, CellularComponent, Compound, Disease, Gene, PharamacologicalClass, SideEffect, and Symptom) all use standardized terminologies and were derived from five different ontologies. The sources and counts of each node and edge type are detailed in Supplementary Tables 1A,B.

### Connecting EHRs To SPOKE

EHRs were connected to SPOKE *Disease, Symptom, SideEffect, Compound*, and *Gene* nodes. To connect to *Disease* nodes, ICD9/10 (Steindel 2010) codes in the EHRs were translated to *Disease* Ontology identifiers (Schriml et al., 2012; Kibbe et al., 2015). Since this relationship was used to select the patient cohort, we manually curated the mappings. The connection to *Symptom* and *SideEffect* nodes was also made from translating the ICD9/10 codes via MeSH identifiers and CUI respectively. The relationship between *Compound* nodes and EHRs was derived by mapping RxNorm to the FDA-SRS UNIIs (Unique Ingredient Identifiers) to DrugBank Identifiers. Lab tests were connected to multiple node types in SPOKE using the Unified Medical Language System (UMLS) Metathesaurus (Bodenreider, 2004). The LOINC (McDonald et al., 2003) codes in the EHRs were mapped to CUI and then mapped to a second CUI (CUI2) using UMLS relationships. A connection between LOINC and SPOKE would be made if CUI2 could be translated to a node in SPOKE. CUIs with nonspecific relationships were excluded. There are 70,843 unique codes found in the Diagnosis, Medication Orders, and Labs tables in the UCSF EHRs, 70,842 of which mapped to 3,527 nodes in SPOKE. Of these, 3,233 were seen in the complex disease cohort and were used as the SPOKE Entry Points (SEPs).

### Generating Propagated SPOKE Entry Vectors

First, we initialized a n x n SEP transition matrix (where n = the number of SEPs) and set every value to zero. Then for each patient in the complex disease cohort, we created a binary vector of the SEPs in their EHRs and divided it by the sum of the vector. This patient vector was then added to the rows of the SEP transition matrix that corresponded to the SEPs found in the patient’s EHRs. Once every patient was accounted for, the SEP transition matrix was transposed and divided by the sum of the columns.

Next, we made an adjacency matrix using the edges in SPOKE to create a SPOKE transition probability matrix (TPM) in which each column sums to 1. The SPOKE TPM was then multiplied by 1-β where β equals the probability of random jump. An extra row was then added to the SPOKE TPM and filled with β.

Last, the Propagated SPOKE Entry Vectors (PSEVs) were generated using a modified version of the PageRank algorithm (Page et al., 1999; Haveliwala 2002). In this version of PageRank, for each PSEV, the random walker traverses the edges of SPOKE until randomly jumping out of SPOKE (at probability β) to the given SEP. The walker will then enter back into SPOKE through any SEP using the probabilities found in the corresponding column of the SEP transition matrix. The walker will continue this cycle until the difference between the rank vector in the current cycle and the previous cycle is less than or equal to a threshold (α). The final rank vector is the PSEV and contains a value for every node in SPOKE that is equivalent to the amount of time the walker spent on each given node.

### BMI GWAS

Genes were selected from the GWAS Catalog if they were associated with an increase in BMI and were genome wide significant.

### Disease Benchmark

#### Generating *Disease* PSEV matrix for benchmark

We created *Disease* benchmark PSEV matrix (PSEV^ΔDD, ΔDG^) by removing the *Disease*-*Disease* and *Disease*-*Gene* relationships in SPOKE prior to PSEV creation. We then used z-scores to normalize the PSEV^ΔDD, ΔDG^ and ranked the elements for each type of node.

#### Random *Disease* matrix

In order to test the importance of the edges between SEPs and SPOKE as well as SPOKE’s internal edges, we generated three types of random PSEVs. First, we created a completely random PSEV matrix by using the Fisher–Yates method to permute the SPOKE nodes for each *Disease* PSEV (PSEV^random^). Second, for each edge type in SPOKE, we randomly shuffled the edges prior to PSEV creation (PSEV^shuffled_SPOKE^). Third, we shuffled the edges between the SEPs and SPOKE prior to PSEV creation (PSEV^shuffled_SEP^). It should be noted that when creating PSEV^shuffled_SEP^, all SPOKE relationships were maintained. Additionally, SEP-SPOKE edges were only shuffled once and therefore any relationships coming directly from the merged EHRs to the SEPs would be conserved. Once random PSEVs were created they were normalized using z-scores

#### Inferring *Disease*-*Gene* Relationships From PSEVs

In addition to looking at *Disease*-*Disease* relationships, we examined the ability of PSEVs to rank the *Disease*-ASSOCIATES_DaG-*Gene* relationships from SPOKE. The Disease-ASSOCIATES_DaG-*Gene* edges (n=12,623) in SPOKE come from four sources: the GWAS Catalog (MacArthur et al., 2017), DISEASES (Pletscher-Frankild et al., 2015), DisGeNET (Pin ero et al., 2015; Pin ero et al., 2016), and DOAF (Xu et al., 2012).

After z-score normalizing the PSEV matrix, within each Disease *PSEV, Gene*s were ranked 1 to 5,392 or 20,945 when using only *Genes* that are associated with at least one *Disease* or the full set of *Genes* accordingly, such that a *Gene* ranked 1 would denote the most important *Gene* for a given *Disease* based on the PSEV matrix. Then for each *Disease* PSEV, K Genes were selected where K was equal to the number of *Genes* are associated with a given *Disease*. The p-values for ability of each *Disease* PSEV to correctly rank the associated *Genes* were then combined using Fisher’s method (Fisher 1992). This evaluation was applied to the original PSEV, benchmark PSEV, and all three random networks (Figure 4A-B).

#### Creating Disease-Gene heat map

The PSEV^ΔDD, ΔDG^ matrix was filtered such that it only contained Disease PSEVs and the Gene elements that are associated with at least one Disease in SPOKE (m=137, n=5,392). This was then used as input into the seaborn clustermap package in python with the settings method=’average’ and metric=’euclidean’.

### Shortest paths between SEP to target nodes

To understand how the PSEVs were able to recover deleted relationships we traced from the target node back to the contributions of each SEP. To achieve this, we z-score normalized the original SEP transition matrix used to calculate the PSEVs. Then we created a SPOKE only PSEV matrix (PSEV^SPOKE-only^) that forces the random walker to randomly restart (B=0.33) from a single SEP. The PSEV^SPOKE-only^ matrix was create using SPOKE with deleted *Disease-Disease* and *Disease-Gene* edges or *Compound-Compound* and *Compound-Gene* edges when recovering the paths for PSEV^ΔDD, ΔDG^ and PSEV^ΔCC, ΔCG^ accordingly. The PSEV^SPOKE-only^ matrix allows to identify the contribution of an individual SEP to any of the downstream nodes. We then took the product of a given *Disease* or *Compound* transposed vector from the SEP transition matrix with the PSEV^SPOKE-only^ to generate contributions of each SEP to the target node. The most important SEP were selected if they were in the top 0.1 percentile of contributors. We then found the shortest paths between the important SEPs and the target node.

## SUPPLEMENTARY TEXT

### Inferring *Disease*-*Disease* Relationships From PSEVs

Utilizing the normalized original matrix (PSEV), benchmark matrix (PSEV^ΔDD, ΔDG^) and the three random PSEV matrices, we checked to see if the deleted SPOKE *Disease*-RESEMBLES_DrD-*Disease* edges could be inferred directly from the PSEV matrices. The *Disease*-RESEMBLES_DrD-*Disease* edges in SPOKE were derived using MEDLINE co-occurrences (n=1,086). This evaluation mirrors that used to test the recovered *Disease-Gene* relationships. However, in this case the *Disease*s elements (n=129 using Diseases that resemble at least one other Disease or n=137 for entire set of Diseases in SPOKE) in each *Disease* PSEV were ranked such that the one ranked 1 would denote the most similar to a given *Disease*. All PSEV matrices were evaluated using this method (Supplementary Figure 2).

### Recovering Deleted *Disease* Resembles *Disease* Relationships

We next used PSEV to create a *Disease*-*Disease* network (DD^PSEV^) as we did the *Disease*-*Gene* networks and used a similar strategy to build background networks as comparators (DDPSEVΔDD, ΔDG, DDRANDOM, DDSPOKE SHUFFLE, and DDSEP SHUFFLE) using the original, benchmark and three random PSEV matrices. These *Disease*-*Disease* networks were then evaluated by the number of edges they shared with the *Disease*-RESEMBLES_ *Disease*_(DrD)-network from SPOKE (DD^SPOKE^). The RESEMBLES_DrD edges in SPOKE were created using the most statistically significant MEDLINE term co-occurrences (n=1,086, p<0.005; Himmelstein et al. 2017). Again, we found that DD^PSEV^ (and even DD^PSEV^^ΔDD, ΔDG^) was able to recover more of the deleted edges (on average 4.7x and 3.7x accordingly) than any of the three random networks (Supplementary Figure 2B).

Interestingly, one of the three random networks (DD^SPOKE SHUFFLE^) performed significantly better than the other two. We hypothesize this is due to the fact that some *Disease*-*Disease* relationships are observable in the EHRs as co-morbidities and misdiagnoses. This information is then feed directly into the *Disease* SEPs, making *Disease*s that resemble other *Disease*s significant in the PSEVs. Since this relationship does not always need to traverse paths in SPOKE, it is observable in the DD^SPOKE SHUFFLE^. In contrast, in DD^SEP SHUFFLE^ the altered mappings between the SEPs and SPOKE disrupt observable relationships in the EHRs, which in turn inhibits the prioritization of *Disease* nodes. These results highlight the accuracy of the mappings between EHR concepts to nodes in SPOKE.

Additionally, in order to learn how we are able to correctly identify related *Diseases* even after deleting *Disease-Gene* and *Disease-Disease* edges from SPOKE, we retraced the shortest paths between significant SEPs of a given *Disease* to its target related *Disease(s)*. Figure 2A shows how Hypertension was ranked as a top *Disease* in the Type 2 Diabetes PSEV^ΔDD, ΔDG^. The “pressure” from the EHRs of Type 2 Diabetes patients pushes the flow of information to the *Anatomy* in which Hypertension is localized, *Symptoms* presented by Hypertension, and *Compounds* that treat or palliate Hypertension. This flow of information makes Hypertension a top ranked *Disease* for Type 2 Diabetes. Further, this pattern of information flow, particularly through *Anatomy* and *Symptom* nodes, is very conserved in the shortest paths between *Disease* pairs.

### Compound Benchmark

#### *Compound*-*Compound* PSEV Based Network

We created *Compound* benchmark PSEVs (PSEV^ΔCC, ΔCG^) by removing the *Compound*-*Compound* and *Compound*-*Gene* relationships in SPOKE prior to PSEV creation. We then used z-scores to normalize the PSEV^ΔCC, ΔCG^.

#### Random *Compound* PSEVs

The three random *Compound* PSEV matrices were derived in the same way as the random *Disease* PSEV matrices. First, PSEV^RANDOM^ was created by permuting the nodes in the *Compound* PSEVs using the Fisher–Yates method. Second, PSEV^SPOKE Shuffle^ was created by shuffling the edges within SPOKE, by edge type. Third, PSEV^SEP Shuffle^ was created by shuffling the edges between SEPs and SPOKE, by edge type. Neither *Compound*-*Compound* or *Compound*-*Gene* edges were deleted prior to random PSEV calculation. All random PSEV matrices were then z-score normalized.

#### Inferring *Compound-Protein* binding partners using EHR embeddings

Employing the original matrix (PSEV), benchmark matrix (PSEV^ΔCC, ΔCG^) and three random matrices (PSEV^random^, PSEV^shuffled_SPOKE^, and PSEV^shuffled_SEP^) we tested whether the molecular targets of a given compound were ranked higher in that *Compound*’s PSEV. To test this we used the *Compound*-BINDS_CbG-*Gene* edges in SPOKE which were derived from a *Compound*’s protein targets from BindingDB (Chen et al., 2001; Gilson et al., 2016), DrugBank (Law et al., 2014; Wishart et al., 2006), and DrugCentral (Ursu et al., 2017) (11,571 edges).

Though this method of evaluation is very similar to the previous methods, it differed in that we selected a fixed number of top K ranked nodes to select from each *Compound* PSEV (K=150). The decision to choose a fixed K was based on the fact that the average number of Gene binding partners per Compound was much smaller than the average number of Genes that associate with Diseases. The value of K was calculated by finding the point at which the patient population no longer contributes positively to the rank of the target *Gene*. The simplest way to calculate patient contribution to the target *Gene* is through proportion of patients on a given *Compound* that have been diagnosed with a *Disease* that is related to the target Gene (Supplementary Fig 3C). This is computed by z-score normalizing the transition probability matrix and summing the values of *Diseases* that are related to the target *Gene* for a given *Compound*. We then plot the aggregated z-scores against rank to find the point in which the aggregated z-scores reaches zero (K=150; Supplementary Fig 3C).

Interestingly, we found that the most significant negative information flow (right end of the plot) was associated with the worst ranked *Genes* and often corresponded to contraindications. For example, Tolmetin, a non-steroidal anti-inflammatory drug, targets *PTGS1* -a gene known to be related to hypertension (Radi, Z., et al. 2007; Bruno, A., et al. 2014; Supplementary Fig 3A). However, Tolmetin is contraindicated for hypertension because it increases the risk of adverse cardiovascular events. As a result, within the population of patients that were prescribed Tolmetin, the number of patients that were also diagnosed with hypertension was fewer than expected. This causes negative information flow through *PTGS1* in the Tolmetin PSEV.

Next, selecting the top 150 *Genes* per *Compound PSEV*, we built *Compound*-*Gene* networks using the original (CG^PSEV^), benchmark (CG^PSEVΔCC, ΔCG^), and three random PSEV matrices (CG^RANDOM^, CG^SPOKE SHUFFLE^, and CG^SEP SHUFFLE^) respectively. We then compared the number of overlapping edges between the CG^SPOKE^, a *Compound-Gene* network created with the *Compound*-BINDS_CbG-*Gene* edges in SPOKE, and the other CG networks. When selecting the top K *Genes* using only *Genes* that have at least one BINDS_DbG edge, we found that CG^PSEVΔCC, ΔCG^ and CG^PSEV^ shared on average 1.9x and 6.9x more edges than the three random networks (Supplementary Fig. 3B) and when selecting the top K from all Gene nodes in SPOKE, the sharing increased to 4.3x and 51.5x respectively (Supplementary Fig. 3B insert). These results show that adding patient information from the EHRs enables the discovery of Compound-Gene relationships in SPOKE.

Finally, to unravel how *Compound* binding partners are highly ranked in PSEVs even after *Compound-Gene* and *Compound-Compound* edges are deleted, we again retraced the shortest paths between significant SEPs and the target *Gene*. Ursodeoxycholic acid is a cholesterol-lowering medication that can also be used to dissolve gallstones and treat liver disorders and is known to target the protein ABCB11, a member of the superfamily of ATP-binding cassette (ABC) transporters (Green et al., 2000; Schuetz et al., 2001; Mita et al., 2005). Supplementary Figure 3A shows how EHRs from patients prescribed Ursodeoxycholic acid guide the flow of information to ABCB11. The information is driven towards *BiologicalProcess* and *Pathway* nodes that ABCB11 participates in and *Diseases* that are localized in *Anatomies* that ABCB11 is expressed or regulated in. Since *Gene* nodes only represent a small fraction of SEPs, this pattern of flow from SEP to target *Gene* is not very common because it includes a *Gene* node (gamma-glutamyltransferase 1, *GGT*) as one of the SEPs. High levels of GGT are often associated with liver or bile duct diseases, which explains why patients may benefit from this drug, as well as informs the connection to ABCB11. More commonly, the shortest paths will involve information flow through *Disease, Anatomy*, and occasionally *Gene* nodes.

#### *Compound* fingerprint similarity in EHR embeddings

Analogous to generating the *Disease-Disease* networks, we created *Compound*-*Compound* networks using the top K ranked *Compound* nodes in the original (CC^PSEV^), benchmark (CC^PSEVΔCC,ΔCG^), or random PSEV (CC^RANDOM^, CC^SPOKE SHUFFLED^, and CC^SEP^^SHUFFLED^) matrices, where K equals the number of similar Compounds to a selected Compound. Then we created a fingerprint-based *Compound*-*Compound* network (CC^SPOKE^) using the *Compound*-RESEMBLES_CrC-*Compound* edges (n=7,703) in SPOKE. The *Compound*-RESEMBLES_CrC-*Compound* edges in SPOKE were derived using the similarity between two Compounds extended connectivity fingerprints (Rogers and Hahn, 2010; Morgan, 1965) and filtered based on their Dice coefficient (Dice, 1945; Himmelstein et al. 2017). Next, we computed the number of edges that were shared between CC^SPOKE^ and the other *Compound*-*Compound* networks. We found that the observed number of shared edges in CC^PSEV^^ΔCC, ΔCG^ and CC^PSEV^ were on average significantly higher than random (4.4x and 15.2x) when selecting from the set of Compounds that resembles at least one other Compound and even higher (4.9x and 17.6x) when selecting from the entire set of nodes in SPOKE (Supplementary Figure 4B). Again the p-values in the figure were calculated using Fisher’s method to combine the p-values for selecting the top K *Compounds* from each *Compound* PSEV^ΔCC, ΔCG^.

Just as when we inferred *Disease*-*Disease* relationships, we noticed that CC^SPOKE SHUFFLED^ performed better than the other two random networks. Again, this is likely because we attempted to predict relationships that can sometimes be observed without traversing SPOKE because they are observable in the EHRs. Therefore, shuffling the edges within SPOKE won’t greatly impact this prediction. Furthermore, these results also demonstrate that we are correctly mapping medication orders in the EHRs to *Compound* nodes in SPOKE.

To elucidate how the benchmark PSEVs could infer whether two compounds were similar, we again found the shortest paths between the important SEPs and target (*Compound*) node. We found that in order to connect Compounds, the random walker usually followed one of two path patterns. In one pattern, the information from the patient population on a given *Compound* is “pushed” through shared *SideEffects* and *PharmacologicalClasses*. For example, Tioconazole resembles Sertaconazole (similarity=0.80) and in order to connect the two Compounds pressure from patients on Tioconazole must move information flow through the *SideEffects* Pruritus, Erythema, Dry skin, and Application site reaction and the *PharmacologicalClass* Azoles (Supplementary Fig. 4A left). The other shortest path pattern for recovering similar *Compounds* is observed when two *Compounds* treat the same *Disease*. An example of this is seen when connecting Trihexyphenidyl to Procyclidine (similarity=0.98; Supplementary Fig. 4A right) which both are used to treat Parkinson’s disease (PD). Here, most of the weight from the EHRs of patients on Trihexyphenidyl is coming from PD and nodes related to PD: Trihexyphenidyl (*Compound* treats PD), Dyskinesias (*Symptom* presented by PD), and Tremor (*Symptom* presented by PD). This results in significant information flow to the Procyclidine node. These results prove the PSEVs ability to identify Compounds with similar structures as well as illustrate what components of the EHRs and relationships of SPOKE are most critical to inform that decision.

### SideEffect to Anatomy Benchmark

#### MEDLINE Co-occurrence Gold Standard

MEDLINE yearly publishes the co-occurrences of MeSH terms found on Pubmed publications. After converting *Anatomy* and *SideEffect* identifiers to MeSH IDs we created a counts matrix for co-occurring *Anatomy* and *SideEffect* terms. Out of the 699,745 possible pairs, 222,224 had at least one co-occurrence). Then we preformed χ^2^ to determine the significance of the *Anatomy*-*SideEffect* MEDLINE relationships. Since 51% of relationships had a p-value less than or equal to 0.05, we decided to strengthen the filter to the top 5% of p-values (p=7.4E-75) leaving 11,112 *Anatomy*-*SideEffect* pairs.

#### PSEV Benchmark *Anatomy*-*SideEffect* Network

First, we used z-score to normalize the PSEV matrix. Then we transposed the PSEV matrix (PSEV^T^) to obtain a vector (n=3,233) for every node in SPOKE. This vector describes the importance of a given SPOKE node for each SPOKE Entry Point (SEPs). Next, vectors from PSEV^T^ were then used to calculate the cosine similarity between *Anatomy* and *SideEffect* nodes. Finally, the similarities were ranked (1 to 699,745), such that a rank of 1 signified the most similar *Anatomy*-*SideEffect* pair in the matrix.

#### Random *Anatomy*-*SideEffect* Networks

To create a random PSEV^T^ matrix, the normalized benchmark PSEV^T^ was shuffled using the Fisher–Yates method to randomly permute the rows of the matrix. The random PSEV matrix was then used to calculate the cosine similarity between the *Anatomy*-*SideEffect* pairs and ranked from 1 to 699,745 in the same way as the benchmark matrix.

#### Overlapping *Anatomy*-*SideEffect* Links

Benchmark and random *Anatomy*-*SideEffect* networks were created using the top k (k=1 to 699,745, increasing in intervals of 5%) nodes in PSEV and PSEV^RANDOM^ accordingly. Supplementary Figure 5 shows the overlapping counts and fraction between the RP networks and the 11,112 *Anatomy*-*SideEffect* pairs from MEDLINE. Inserts in Supplementary Figures 5A-C focus on k<= 11,112, corresponding the number of *Anatomy*-*SideEffect* pairs from MEDLINE. The highest fold changes 18.1 over random occurred in the top k=1,000 respectively (Supplementary Figure 5C insert).

#### Recovering the major shortest paths between *SideEffect* and *Anatomy* nodes

First, we needed to find the nodes that contributed most weight to the similarity of the S*ideEffect-Anatomy* pair. Since we used cosine similarity, which is equivalent to the dot product of two unit vectors, we simply multiplied the *SideEffect* and *Anatomy* transposed PSEVs and selected the highest 0.1% of nodes. Those nodes are labeled as top contributors in Supplementary Figures 5D-F. We then found the shortest paths between each top contributor node and the target *SideEffect* and *Anatomy* nodes.

#### *SideEffect*-*Anatomy* relationships in embedded EHR concepts match MEDLINE co-occurrences

Although it is natural to draw a connection between drug side effects and the anatomies they affect (e.g. a headache must somehow relate to the brain), *SideEffect* and *Anatomy* nodes are not directly connected in SPOKE. In fact, in order to get from a *SideEffect* to an *Anatomy* node one must traverse a minimum of three edges. As a result, correctly inferring the relationships between *Anatomy* and *SideEffect* nodes would show that appropriate weights are assigned to distant nodes in the network. To test this, we created a gold standard *SideEffect*-*Anatomy* network using only highly significant relationships from MEDLINE co-occurrences (SeA^MEDLINE^) (p=7.4e-75; n=11,112; avg

Anatomy per SideEffect). Next, we computed a *SideEffect*-*Anatomy* cosine similarity matrix using the transposed PSEV matrix (See methods). We then selected the most similar *SideEffect*-*Anatomy* pairs to create a PSEV-based *SideEffect*-*Anatomy* network (SeA^PSEV^). These relationships were also tested against a random network (SeA^RANDOM^) that was generated by permuting each PSEV, as in the DD^RANDOM^ networks (Supplementary Figure 5).

In the first interval (k=1000), we observed 18.1 times more overlapping edges than expected by chance (Supplementary Figure 5C insert; binomial p value = 9.7E-251). By accurately ranking the relationships between *SideEffect* and *Anatomy* nodes, we further demonstrate that PSEVs are a valid strategy to infer missing links in SPOKE. This result is even more consequential given that *SideEffect* and *Anatomy* nodes are far away in SPOKE.

Similar to before when we found the shortest paths between SEPs and the target node to understand how deleted edges where recovered, we wanted to find the paths that enabled us to learn relationships between *SideEffect* and *Anatomy* nodes. To achieve this, we found the nodes in the transposed PSEVs that contributed the most to the *SideEffect* and *Anatomy* similarity. We then looked at the shortest paths between those nodes and the target *SideEffect* and *Anatomy* nodes. Supplementary Figures 5D-F show examples of these paths. The first example shows how Aggression connects to locus coeruleus (LC), a part of the brain that is involved in emotions, arousal, attention, and stress response (Benarroch E., 2009). The nodes that contribute the most to the similarity are *Compounds* and all have the *SideEffect* Aggression. Additionally, those *Compounds* bind or regulate *Genes* expressed or regulated in the LC as well as treat or palliate *Diseases* localized in the LC (Supplementary Fig 5D). Similarly, Supplementary Figure 5E shows the connection between Anxiety (*SideEffect)* and the LC (*Anatomy)*. Interestingly, the shortest paths between Anxiety or Aggression to the LC only share three nodes: alcohol dependence, epilepsy syndrome, and hypertension. The final example shows the connections between fetal heart rate (*SideEffect)* and the umbilical artery (*Anatomy)* (Supplementary Fig. 5F). This connection is centered on a set of genes that are associated or regulated by Diseases localized in umbilical artery. Those same *Genes* are also targets of or regulated by *Compounds* that impact fetal heart rate. These examples further show that PSEVs can be used to find related biomedical entities and further our understanding of how and why they are connected.

## ACKNOWLEDGEMENTS

We thank Sourav Bandyopadhyay, Riley Bove, Jeffrey Gelfand, Sharat Israni, and Keith Yamamoto for helpful discussions. Partial support for this work was provided by grants from Genentech to A.J.B () and S.E.B (G-54860). The sponsor had no role in the design or implementation of this study. Additionally, we would like to thank Achievement Rewards for College Scientists (ARCS) Scholarship and the NHI BMI Training Grant (T32 GM067547/ 4T32GM067547-14). SEB holds the Heidrich Family and Friends Endowed Chair of Neurology at UCSF.

## SUPPLEMENTARY FIGURE LEGENDS AND TABLES

**Supplementary Figure 1. PSEVs embed first neighbors in SPOKE and learn new relationships.** Imagine the SPOKE network as a set of water pipes and the SEPs as input valves. Pressure from the patient population determines how much water can flow through the valves. The water can then reach downstream SPOKE nodes. The amount of water that flows through each SPOKE node will be specific to the selected patient population. (A) Distribution of ranks in PSEV vectors for first neighbors (blue) and non-first neighbors (red). (B) Multiple sclerosis first neighbors that overlap with top PSEV rank (blue edges) or not in top PSEV rank (red). (C) The top 10 ranked nodes in the PSEV for each node types that don’t directly connect to Multiple sclerosis Disease node in SPOKE (dashed edges)

**Supplementary Figure 2. Recovering deleted *Disease-Disease* edges. (A)** shows how the deleted *Disease-Disease* edge between Type 2 Diabetes and Hypertension is recovered using the pressure generated from the Type 2 Diabetes patients. **(B)** The gold standard *Disease-Disease* network was made from the deleted edges in SPOKE. Plots show the number of *Disease-Disease* relationships using each of the PSEV matrices that overlap with the gold standard network. The pink distributions show the results from the permuted PSEV matrices (PSEV^Random^; 1000 iterations) while the arrows show the results from the original PSEV (blue), PSEV^ΔDD, ΔDG^ (green), PSEV^SPOKE SHUFFLED^ (red), and PSEV^SEP SHUFFLED^ (orange). **(B)** The top K *Diseases* where selected from the set of *Diseases* in the gold standard network or **(B insert)** the entire set of *Disease* in SPOKE. **(F)** The top K *Diseases* where selected from the set of *Diseases* in the gold standard network or **(F insert)** the entire set of *Disease* in SPOKE.

**Supplementary Figure 3. Recovering deleted *Compound-Gene* edges.** Prior to PSEV^ΔCC, ΔCG^ calculation all of the *Compound -Gene* and *Compound -Compound* edges were deleted from SPOKE. It is possible to retrace how PSEV can recover deleted edges (outlined in Figure 4C). **(A)** Shortest paths between the top SEPs of Tolmetin, a non-steroidal anti-inflammatory drug, to its target *PTGS1*. **(B)** The gold standard *Compound-Gene* network was made from the deleted edges in SPOKE (*Compound*-BINDS_CbG-*Gene*). Plots show the number of *Compound-Gene* relationships using each of the PSEV that overlap with the gold standard networks. The pink distributions show the results from the permuted PSEV matrices (PSEV^Random^; 1000 iterations) while the arrows show the results from the original PSEV (blue), PSEV^ΔCC, ΔCG^ (green), PSEV^SPOKE SHUFFLED^(red), and PSEV^SEP SHUFFLED^ (orange). **(B)** The top K *Genes* where selected from the set of *Genes* in the gold standard network or **(B insert)** the entire set of *Gene* nodes in SPOKE. **(C-E)** Determining K threshold for recovering *Compound-Gene* edges. **(C)** The top factor in determining missing *Compound-Gene* edges is whether patients that are on a given compound are also diagnosed with a Disease that is a associated with the target gene. **(D)** Shows the number of recovered *Compound-Gene* relationships at each rank (where 1=top ranked and 1451 is the worst ranked *Gene*). **(E)** Shows how much the patients that are prescribed a given *Compound* are contributing to the rank of the binding partner (missing *Compound-Gene* relationship) of that *Compound* using the flow of information through Diseases as in A. Genes ranked greater than ∼150 are no longer receiving positive patient contribution.

**Supplementary Figure 4. Recovering deleted *Compound-Compound* edges. (A)** Retracing shortest between similar *Compounds*. The paths between Tioconazole to Sertaconazole and Trihexyphenidyl to Procyclidine show two different routes in finding similar compounds. **(B)** The gold standard *Compound-Compound* network was made from the deleted edges in SPOKE (*Compound*-RESEMBLES_CrC-*Compound*). **(B)** The top K *Compound* where selected from the set of *Compound* in the gold standard network or **(B insert)** the entire set of *Compound* in SPOKE.

**Figure 5 MEDLINE Anatomy-SideEffect Relationships are Top Ranked Nodes in PSEV**. Fraction **(A),** count **(B)**, and fold change **(C)** of overlapping edges MEDLINE Anatomy-SideEffect network and PSEV Anatomy-SideEffect network (blue) or random PSEV Anatomy-SideEffect network (red) for different thresholds of PSEV disease similarity. A-C Are shown in 5% similarity intervals of ranked nodes starting with the most similar 5% left and all nodes (100%) right. The inserts in A-C focus on the top 0.14-1.6% of ranked nodes. D-F Examples shortest paths connecting the nodes that contribute the most to the SideEffect-Anatomy similarity to the target SideEffect and Anatomy nodes.

**Supplementary Table 1. SPOKE nodes and edges**. (A) Source(s) and counts of each node type in SPOKE. (B) Source(s) and counts of each edge label in SPOKE.

