## Supplementary Figure 1 for "Integrating Biomedical Research and Electronic Health Records to Create Knowledge Based Biologically Meaningful Machine Readable Embeddings"

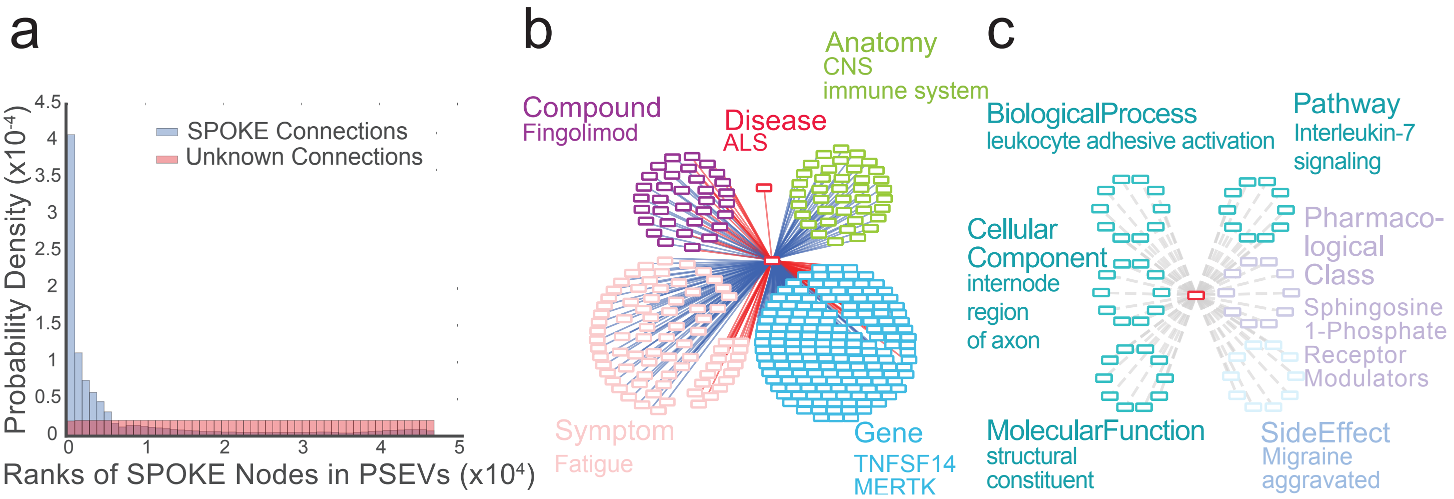


**Supplementary Figure 1. PSEVs embed first neighbors in SPOKE and learn new relationships.** Imagine the SPOKE network as a set of water pipes and the SEPs as input valves. Pressure from the patient population determines how much water can flow through the valves. The water can then reach downstream SPOKE nodes. The amount of water that flows through each SPOKE node will be specific to the selected patient population. **(A)** Distribution of ranks in PSEV vectors for first neighbors (blue) and non-first neighbors (red). **(B)** Multiple sclerosis first neighbors that overlap with top PSEV rank (blue edges) or not in top PSEV rank (red). **(C)** The top 10 ranked nodes in the PSEV for each node types that don't directly connect to Multiple sclerosis Disease node in SPOKE (dashed edges)
