## Supplementary Figure 2 for "Integrating Biomedical Research and Electronic Health Records to Create Knowledge Based Biologically Meaningful Machine Readable Embeddings"

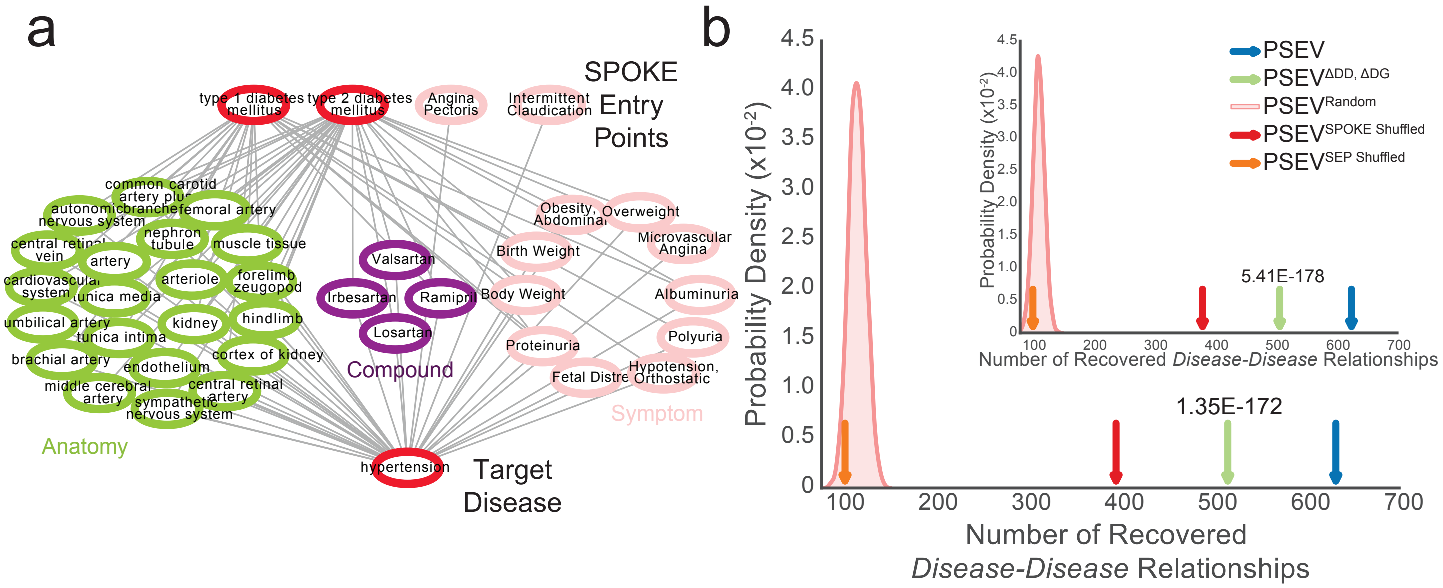


**Supplementary Figure 2. Recovering deleted *Disease-Disease* edges. (a)** shows how the deleted *Disease-Disease* edge between Type 2 Diabetes and Hypertension is recovered using the pressure generated from the Type 2 Diabetes patients. **(b)** The gold standard *Disease-Disease* network was made from the deleted edges in SPOKE. Plots show the number of *Disease-Disease* relationships using each of the PSEV matrices that overlap with the gold standard network. The pink distributions show the results from the permuted PSEV matrices ($PSEV^{Random}$; 1000 iterations) while the arrows show the results from the original PSEV (blue), $PSEV^{\Delta DD, \Delta DG}$ (green), $PSEV^{SPOKE\ SHUFFLED}$ (red), and $PSEV^{SEP\ SHUFFLED}$ (orange). **(b)** The top K *Diseases* where selected from the set of *Diseases* in the gold standard network or (b insert) the entire set of *Disease* in SPOKE.
