## Supplementary Figure 3 for "Integrating Biomedical Research and Electronic Health Records to Create Knowledge Based Biologically Meaningful Machine Readable Embeddings"

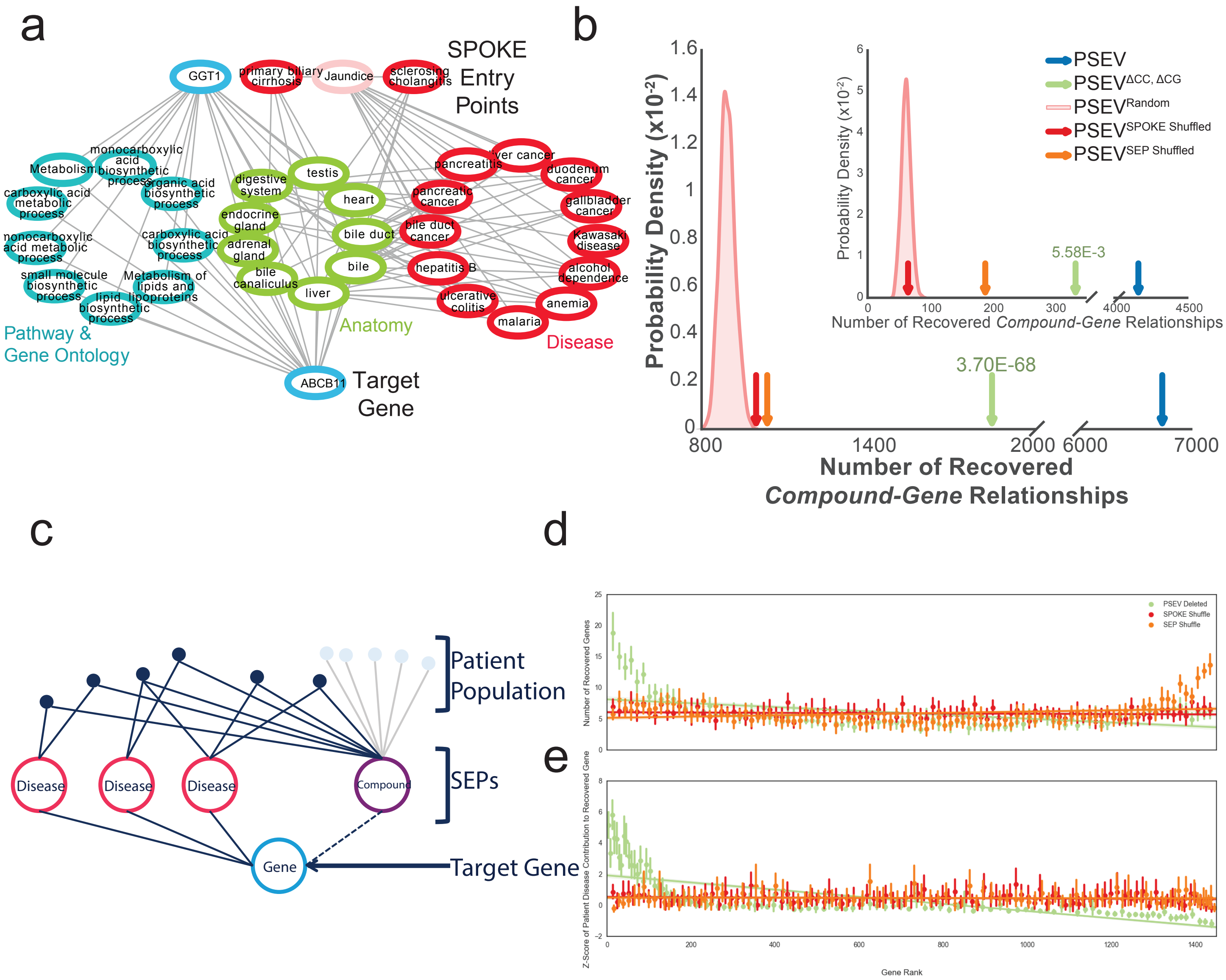


**Supplementary Figure 3. Recovering deleted *Compound-Gene* edges.** Prior to PSEV$^{\Delta CC, \Delta CG}$ calculation all of the *Compound-Gene* and *Compound-Compound* edges were deleted from SPOKE. It is possible to retrace how PSEV can recover deleted edges (outlined in Figure 4C). **(A)** Shortest paths between the top SEPs of Tolmetin, a non-steroidal anti-inflammatory drug, to its target PTGS1. **(B)** The gold standard *Compound-Gene* network was made from the deleted edges in SPOKE (*Compound*-BINDS_CbG-*Gene*). Plots show the number of *Compound-Gene* relationships using each of the PSEV that overlap with the gold standard networks. The pink distributions show the results from the permuted PSEV matrices (PSEV$^{Random}$; 1000 iterations) while the arrows show the results from the original PSEV (blue), PSEV$^{\Delta CC, \Delta CG}$ (green), PSEV$^{SPOKE\ SHUFFLED}$ (red), and PSEV$^{SEP\ SHUFFLED}$ (orange). **(B)** The top K *Genes* where selected from the set of *Genes* in the gold standard network or (B insert) the entire set of *Gene* nodes in SPOKE. **(C-E)** Determining K threshold for recovering *Compound-Gene* edges. **(C)** The top factor in determining missing *Compound*-Gene edges is whether patients that are on a given compound are also diagnosed with a Disease that is a associated with the target gene. **(D)** Shows the number of recovered *Compound-Gene* relationships at each rank (where 1=top ranked and 1451 is the worst ranked *Gene*). **(E)** Shows how much the patients that are prescribed a given *Compound* are contributing to the rank of the binding partner (missing *Compound-Gene* relationship) of that *Compound* using the flow of information through Diseases as in A. *Genes* ranked greater than ~150 are no longer receiving positive patient contribution.
