## Supplementary Figure 4 for "Integrating Biomedical Research and Electronic Health Records to Create Knowledge Based Biologically Meaningful Machine Readable Embeddings"

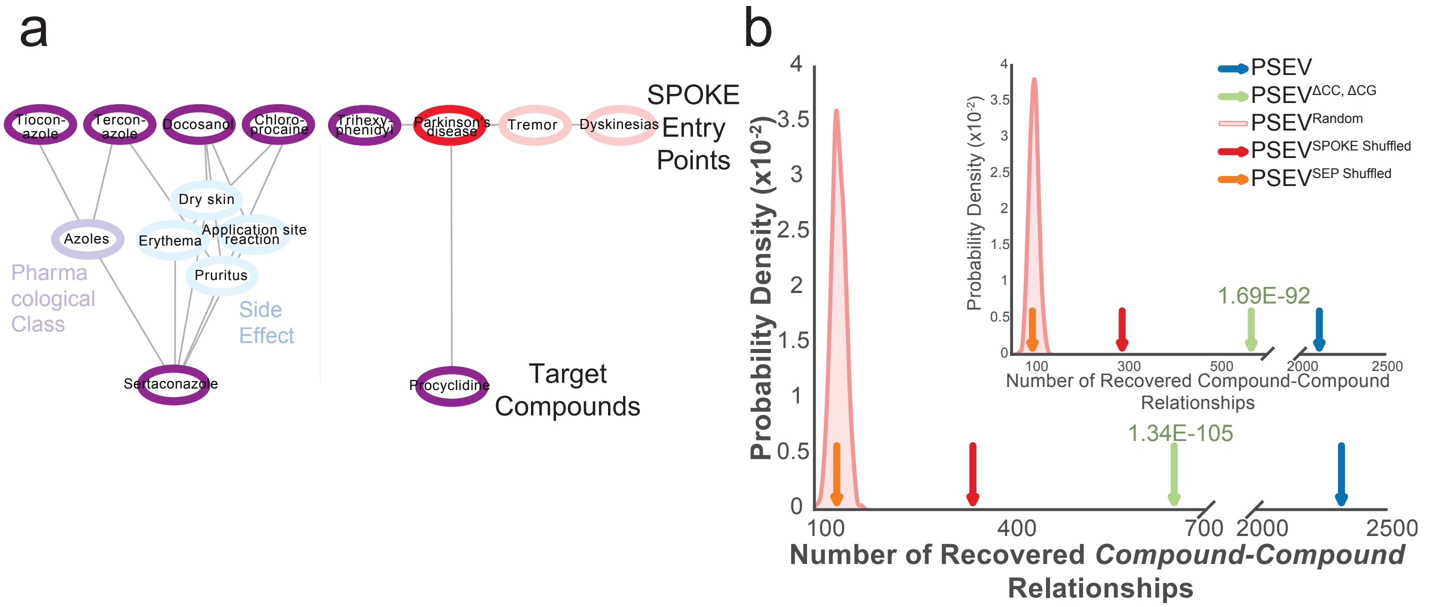


**Supplementary Figure 4. Recovering deleted *Compound-Compound* edges. (a)** Retracing shortest between similar Compounds. The paths between Tioconazole to Sertaconazole and Trihexyphenidyl to Procyclidine show two different routes in finding similar compounds. **(b)** The gold standard *Compound-Compound* network was made from the deleted edges in SPOKE (*Compound*-RESEMBLES_CrC-*Compound*). **(b)** The top K *Compound* where selected from the set of *Compound* in the gold standard network or **(b insert)** the entire set of *Compound* in SPOKE.
