## Supplementary Table 1 for "Integrating Biomedical Research and Electronic Health Records to Create Knowledge Based Biologically Meaningful Machine Readable Embeddings"

| Nodes | | |
|---|---|---|
| Node Name | Source | Count |
| Gene | Entrez Gene | 20945 |
| BiologicalProcess | Gene Ontology | 11381 |
| SideEffect | UMLS via SIDER 4.1 | 5734 |
| MolecularFunction | Gene Ontology | 2884 |
| Compound | DrugBank | 1552 |
| CellularComponent | Gene Ontology | 1391 |
| Pathway | Reactome via Pathway Commons | 1308 |
| Symptom | MeSH | 438 |
| Anatomy | Uberon | 402 |
| PharmacologicClass | FDA via DrugCentral | 345 |
| Pathway | WikiPathways | 294 |
| Pathway | PID via Pathway Commons | 220 |
| Disease | Disease Ontology | 137 |
| Total | | 47031 |

| Edges | | |
|---|---|---|
| Edge Name | Source | Count |
| DOWNREGULATES_AdG | Bgee | 102240 |
| UPREGULATES_AuG | Bgee | 97848 |
| RESEMBLES_CrC | Dice similarity of ECFPs | 6486 |
| INCLUDES_PCiC | DrugCentral | 1029 |
| COVARIES_GcG | ERC | 61690 |
| DOWNREGULATES_CdG | LINCS L1000 | 21102 |
| REGULATES_GrG | LINCS L1000 | 265672 |
| UPREGULATES_CuG | LINCS L1000 | 18756 |
| LOCALIZES_DlA | MEDLINE cooccurrence | 3602 |
| PRESENTS_DpS | MEDLINE cooccurrence | 3357 |
| RESEMBLES_DrD | MEDLINE cooccurrence | 543 |
| PARTICIPATES_GpBP | NCBI gene2go | 559504 |
| PARTICIPATES_GpCC | NCBI gene2go | 73566 |
| PARTICIPATES_GpMF | NCBI gene2go | 97222 |
| PALLIATES_CpD | PharmacotherapyDB | 390 |
| TREATS_CtD | PharmacotherapyDB | 755 |
| PARTICIPATES_GpPW | PID via Pathway Commons | 8154 |
| PARTICIPATES_GpPW | WikiPathways | 12587 |
| PARTICIPATES_GpPW | Reactome via Pathway Commons | 63631 |
| CAUSES_CcSE | SIDER 4.1 | 138944 |
| DOWNREGULATES_DdG | STARGEO | 7623 |
| UPREGULATES_DuG | STARGEO | 7731 |
| ASSOCIATES_DaG | DOAF, GWAS Catalog, DisGeNET, DISEASES | 12623 |
| BINDS_CbG | DrugBank (target), PDSP Ki, PubChem, DrugCentral (I[illegible] | 11571 |
| EXPRESSES_AeG | Bgee, TISSUES | 526407 |
| INTERACTS_GiG | Lit-BM-13, hetio-da, Venkatesan-09, Yu-11, hetio-dag, H[illegible] | 147164 |
| Total | | 2250197 |

**Supplementary Table 1. SPOKE nodes and edges. (A)** Source(s) and counts of each node type in SPOKE. **(B)** Source(s) and counts of each edge label in SPOKE.
